## supplementary figures for "Rapid rebalancing of co-tuned ensemble activity in the auditory cortex"

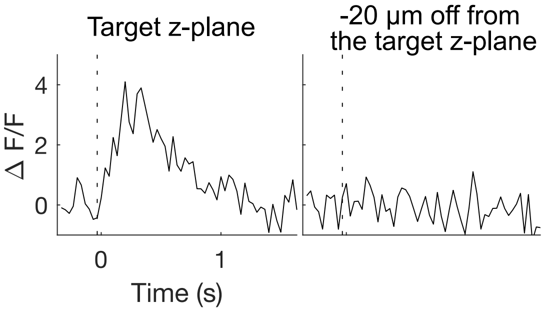


**Supplementary Figure 1. Holographic optogenetic stimulation effectively activates target points.** Holographic optogenetic stimulation effect on five pre-selected cells at the target z-plane (left) and 20 µm off-target z-plane (right; n = 1). No stimulation-driven effect was observed on the off-target cells. Dashed vertical lines indicate the stimulation onset.

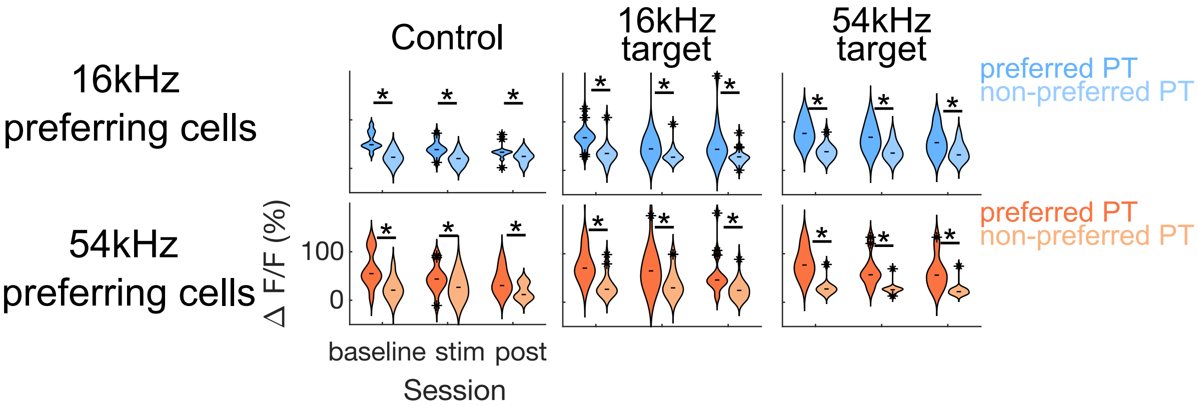


**Supplementary Figure 2. Subgrouped non-target cells show the frequency preference.** Violin plots for average response amplitudes of either 16 kHz preferring cells (blue) or 54 kHz preferring cells (orange) to each frequency for each session and for each condition (either 16kHz 5-cell stimulation, 54kHz 5-cell stimulation, or control with no stimulation at each column). Both groups of cells show greater responses to their preferred frequency (darker shades) regardless of the session (Asterisks above the horizontal line; Mixed-effect model, $p < 0.0001$ for frequency, all $p > 0.05$ across sessions and conditions). Asterisks on the violinplot indicate outlier cells and horizontal lines on the the violinplot indicate medians.

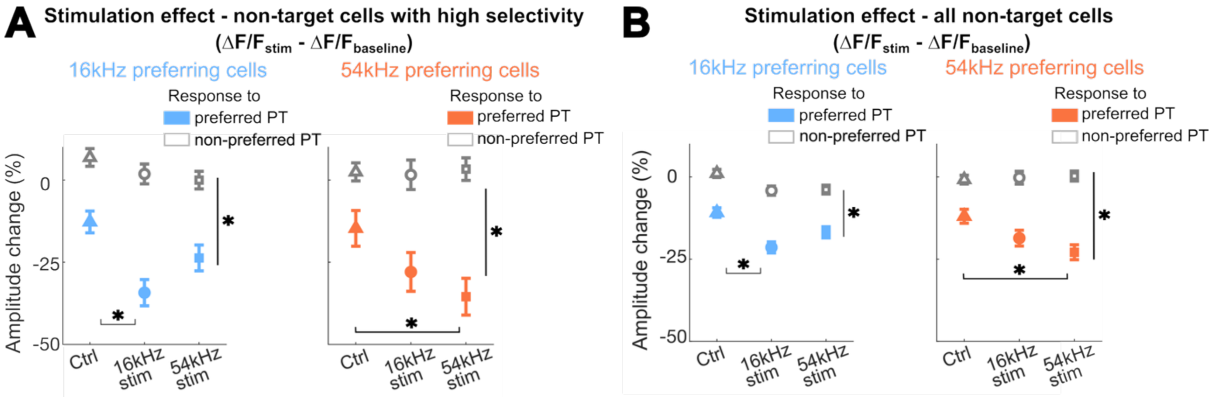


**Supplementary Figure 3. A:** Stimulation effect ($\Delta$ F/F$_{stim}$ - $\Delta$ F/F$_{baseline}$) in 16 kHz (blue) and 54 kHz (orange) preferring cells with high frequency selectivity. Both cell groups show decreased amplitude to their preferred frequency regardless of conditions due to acoustic stimulus-specific adaptation. Only co-tuned cells (16 kHz preferring cells for 16 kHz stimulation or 54 kHz preferring cells for 54 kHz stimulation) show a further decrease in response amplitudes due to the stimulation, when the preferred pure tone (PT) frequency was synchronized. Error bars indicate SEM across cells (*: $p < 0.0001$). **B:** Same as A, but with all frequency selectivity bins pooled. A significant stimulation effect is still observed but attenuated compared to panel A which included only the high frequency selectivity group.
